# Patterns of within-host genetic diversity in SARS-CoV-2

**DOI:** 10.1101/2020.12.23.424229

**Authors:** Gerry Tonkin-Hill, Inigo Martincorena, Roberto Amato, Andrew R J Lawson, Moritz Gerstung, Ian Johnston, David K Jackson, Naomi R Park, Stefanie V Lensing, Michael A Quail, Sónia Gonçalves, Cristina Ariani, Michael Spencer Chapman, William L Hamilton, Luke W Meredith, Grant Hall, Aminu S Jahun, Yasmin Chaudhry, Myra Hosmillo, Malte L Pinckert, Iliana Georgana, Anna Yakovleva, Laura G Caller, Sarah L Caddy, Theresa Feltwell, Fahad A Khokhar, Charlotte J Houldcroft, Martin D Curran, Surendra Parmar, The COVID-19 Genomics UK (COG-UK) Consortium, Alex Alderton, Rachel Nelson, Ewan Harrison, John Sillitoe, Stephen D Bentley, Jeffrey C Barrett, M. Estee Torok, Ian G Goodfellow, Cordelia Langford, Dominic Kwiatkowski, Wellcome Sanger Institute COVID-19 Surveillance Team

**Affiliations:** Wellcome Sanger Institute, Hinxton, United Kingdom; European Bioinformatics Institute, Hinxton, United Kingdom; Department of Medicine, University of Cambridge; Department of Pathology, University of Cambridge; The Francis Crick Institute, London; Cambridge Institute of Therapeutic Immunology and Infectious Disease, University of Cambridge; Public Health England, Cambridge, UK; Nuffield Department of Medicine, University of Oxford

**Keywords:** SARS-CoV-2, within-host, transmission, mutational spectrum

## Abstract

Monitoring the spread of SARS-CoV-2 and reconstructing transmission chains has become a major public health focus for many governments around the world. The modest mutation rate and rapid transmission of SARS-CoV-2 prevents the reconstruction of transmission chains from consensus genome sequences, but within-host genetic diversity could theoretically help identify close contacts. Here we describe the patterns of within-host diversity in 1,181 SARS-CoV-2 samples sequenced to high depth in duplicate. 95% of samples show within-host mutations at detectable allele frequencies. Analyses of the mutational spectra revealed strong strand asymmetries suggestive of damage or RNA editing of the plus strand, rather than replication errors, dominating the accumulation of mutations during the SARS-CoV-2 pandemic. Within and between host diversity show strong purifying selection, particularly against nonsense mutations. Recurrent within-host mutations, many of which coincide with known phylogenetic homoplasies, display a spectrum and patterns of purifying selection more suggestive of mutational hotspots than recombination or convergent evolution. While allele frequencies suggest that most samples result from infection by a single lineage, we identify multiple putative examples of co-infection. Integrating these results into an epidemiological inference framework, we find that while sharing of within-host variants between samples could help the reconstruction of transmission chains, mutational hotspots and rare cases of superinfection can confound these analyses.

## Introduction

The SARS-CoV-2 pandemic has caused global disruption and more than one million deaths (1). Genomic analysis has yielded important insights into the origins and spread of the pandemic, and has been an integral part of efforts to monitor viral transmission in the UK, with over 100,000 viral genomes sequenced as of December 2020 by the COVID-19 Genomics UK Consortium (COG-UK) (2).

For purposes of genomic epidemiology, a consensus genome sequence is derived from each sample but deep sequencing data invariably reveal some level of within-host variation (3) and minor alleles are commonly filtered out prior to phylogenetic analysis (4). It has been suggested that SARS-CoV-2, like SARS-CoV-1 evolves within an infected host as a quasispecies, with many mutations (within-host variants) arising which may be beneficial for the virus (3, 5, 6). Although there have been several previous reports on the within-host diversity of SARS-CoV-2 (3, 7, 8) a number of key questions remain to be resolved. Examples of underexplored questions include understanding the extent of sequencing artefacts among within-host variants, the mutational processes dominating SARS-CoV-2 evolution, the action of selection on within-host variants and the extent to which superinfection or co-transmission of multiple lineages confound within-host diversity analyses (9). Better understanding of these questions can shed light on the evolution of SARS-CoV-2 and could inform efforts to reconstruct transmission chains with genomic data.

To address these questions, we performed Illumina deep sequencing of over a thousand SARS-CoV-2 samples collected in March and April 2020 in the East of England. Two libraries were sequenced for each sample with separate reverse transcription (RT), PCR amplification and library preparation steps in order to evaluate the quality and reproducibility of within-host variant calls. To develop reliable methods for analysing within-host variation in the context of ongoing genomic epidemiological studies, we used the ARTIC protocol which is a common method used for SARS-CoV-2 genome sequencing by many labs around the world (10). Analyses of the data provided insights into the extent of within-host diversity, the patterns of mutagenesis and the extent of selection, and suggested that amplification biases, hypermutable sites and co-infection complicate the use of within-host diversity for epidemiological purposes.

## Results

### Reliable detection of within-host mutations from amplicon sequencing

We generated sequencing data for 1,181 samples in duplicate at a median sequencing coverage ranging from 998 to 49,025 read depth per replicate sample (Supplementary Figure 1; Supplementary Table 1). To study variable sites within an individual, including errors and within-host mutations, for each sample we first identified a ll variable s ites w ith variant a llele f requency *>* 0.5% and at least 5 supporting reads. Comparison of calls between replicates revealed that, while some samples showed highly-concordant variants between replicates, others showed high discordance (Supplementary Figure 2). The discordance between some replicates suggested that a small number of molecules may be disproportionately amplified in some samples, amplifying both RT/PCR errors and rare within-host mutations to high allele frequencies. We quantified the discordance between replicates using a beta-binomial model (Methods) and correlated this with the diagnostic qPCR Ct values, which are inversely related to the number of viral molecules within the samples. Samples with Ct ≥ 24 showed considerable discordance in allele frequencies between replicates (Supplementary Figure 3) but the vast majority of samples with Ct*<* 24 showed good concordance between replicates. This indicated that as the viral loads decrease, amplification b iases a nd a rtefacts a re c ommon a nd c an impact within-host diversity analyses using RT-PCR protocols. For all subsequent analyses we used only those within-host variants that were statistically supported by both replicates (Methods). To reliably detect within-host variants with the ARTIC protocol, we used ShearwaterML, an algorithm designed to detect variants at low allele frequencies using a site-specific error model learned from a collection of reference samples (11, 12) (Methods). Two samples were excluded, as they had an unusually high number of minor variant calls unlikely to be of biological origin, leaving 1,179 samples for analysis, comprising 1,121 infected individuals of whom 49 had multiple samples. Within each sample, we classified variant calls into major variants (≥ 95% allele frequency) and within-host variants (*<* 95%). In total, we identified 18,888 putative variants (Supplementary Table 2), including 7,190 major variants (affecting 948 sites) and 11,698 within-host variants (affecting 5,625 sites). Within-host variants included 7,561 single-nucleotide substitutions, 3,235 small deletions, 548 small insertions and 354 putative multi-nucleotide variants (Methods). The allele frequency spectrum was dominated by fixed mutations that were common to all viral RNA molecules in a sample, and a tail of mostly low frequency variants, as previously described (Lythgoe et al. 2020) (Figure 1A-C). The mean and median allele frequencies of within-host variants was 7.2% and 1.9%, respectively. Overall, within-host variants were detected in the vast majority of samples (95.2%), with a median of 8 within-host variants per sample. This is likely to be an underestimate due to low sensitivity to within-host variants at lower allele frequencies and due to the stringent requirement that a variant is detected as significant in both replicates.

**Fig. 1.**
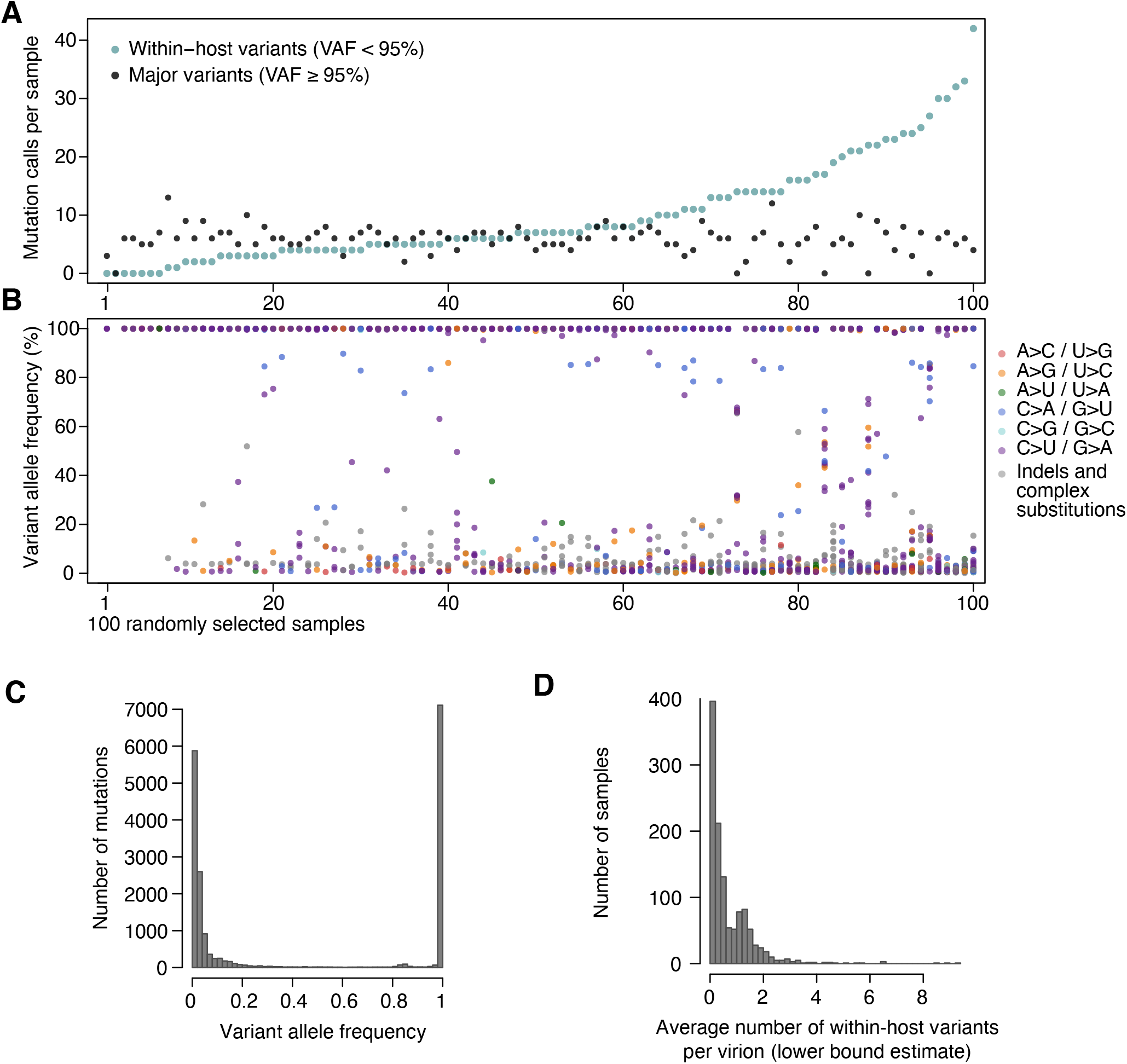
Allele frequencies and mutation burden. (A) Number of variants per sample (y-axis) for a random selection of 100 samples (x-axis). Black and green dots respectively indicate the number of major and within-host variants in each sample. (B) VAF of the variants detected in the 100 samples, coloured by mutation type. (C) Histogram of the VAFs of all mutation calls. Note that variants shared across samples are counted multiple times and that the 7,190 major variants correspond to 961 different changes in 948 different sites. (D) Histogram of the lower bound estimates of the mean number of within-host variants per viral genome per sample.

### The mutational spectra reveal strong strand asymmetries

Analysis of the mutational spectrum can yield insights into the dominant mutational mechanisms underlying the evolution of SARS-CoV-2 during the pandemic. Consistent with previous reports (13), the mutational spectrum of within-host variant closely resembles that of major variants (Figure 2). The spectrum shows two striking features: a dominance of C>U and G>U changes with weak extended sequence context, and a large asymmetry between the plus and minus strands, inferred from the high C>U/G>A and G>U/C>A ratios when mutations are mapped to the reference (plus) strand. C>U mutations account for 47% of all within-host point mutations compared to 5.9% for G>A mutations (C>U mutations in the minus strand), and G>U mutations for 14.5% of mutations compared to 2.2% of C>A mutations (Methods). Normalised for sequence composition, the plus/minus strand ratios of the rates of C>U and G>U are 9.9-fold and 8.2-fold, respectively.

**Fig. 2.**
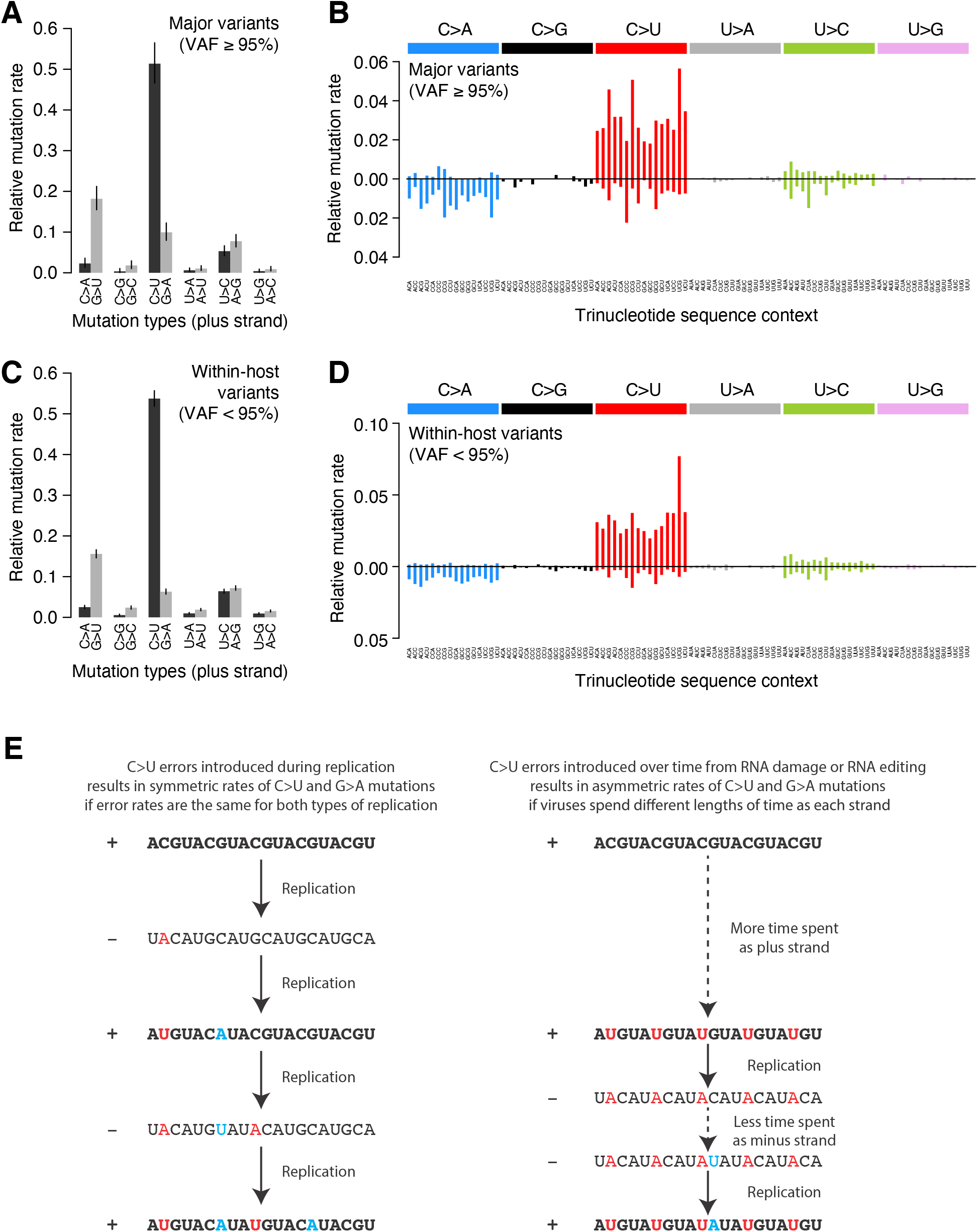
Mutational spectra. (A,C) Mutational spectra (without sequence context) of major (A) and within-host (C) variants, as mapped to the reference strand. Asymmetries suggest different mutation rates in the plus and minus strands. Error bars depict Poisson 95% confidence intervals. (B,D) Mutational spectra in a 96-trinucleotide context of major (B) and within-host (D) variants, as in (27). Mutations are represented as mapped to the pyrimidine base, depicted above the horizontal line if the pyrimidine base is in the reference (plus) strand and below it if the pyrimidine base is in the minus strand. Within-host variants observed across more than one sample can represent a single ancestral event or multiple independent events. To prevent highly-recurrent events from distorting the spectrum, within-host variants observed across multiple samples were counted a maximum of 5 times in (C) and (D). (E) A diagram illustrating how asymmetrical mutation rates of C>U and G>A could be driven by viral sequences spending a longer time as plus strand molecules.

These asymmetries appear difficult to explain if mutations were direct replication errors. Since any given viral RNA molecule has undergone the same number of plus-to-minus and minus-to-plus replication steps, replication errors are only expected to cause these asymmetries if both steps have very different error rates. For example, if the polymerase had a high C>U error rate in both strands, we would expect a symmetric number of C>U and G>A mutations in the plus strand, as C>Us would be introduced at equal rates when copying the plus or the minus strands (Figure 2E). Within a cell, the fact that minus templates are copied many times could theoretically lead to a viral population with fewer G>A mutations but at higher frequencies (14, 15). However, the fact that the same asymmetries are observed for major variants, which typically represent the fixation of a single genotype, makes this an unlikely factor. Thus, unless the error rates of the RNA-dependent RNA-polymerase in SARS-CoV-2 are very different on both strands, which may be possible given its multisubunit structure (16), replication errors seem unlikely to explain the observed asymmetries.

The strong strand asymmetries observed may be more consistent with RNA damage or RNA editing of the plus strand dominating the accumulation of mutations in SARS-CoV-2. The plus strand is the infectious genome, exported from the replication organelles, where minus molecules reside, into the cytoplasm (17), translated, packaged into particles and transmitted between cells and hosts. The plus strand is also present within cells in much larger numbers than the minus strand (14). Thus, the plus strand may be expected to accumulate higher rates of damage or editing, which would manifest as strand asymmetries. If we accept this hypothesis, the dominance of C>U over G>A and G>U over C>A, when mapped to the plus strand, suggests that the dominant forms of RNA editing or damage are C>U and G>U.

RNA-editing enzymes in human cells are able to mutagenise single-stranded DNA and RNA molecules and are known anti-viral mechanisms (18, 19). Two families of RNA-editing enzymes in particular have been speculated to contribute to the mutational spectrum of SARS-CoV-2, APOBEC cytosine deaminase enzymes causing cytosine to uracil transitions and ADAR enzymes causing adenosine-to-inosine changes (A>G/U>C mutations) (19). While we see a high rate of C>U changes in the SARS-CoV-2 spectra, we see much lower rates of A>G/U>C mutations than previously suggested (20). While the mutational spectrum induced by all APOBEC enzymes is not fully understood, the activity of the better-understood APOBEC3A and 3B enzymes in human cancers is distinctly characterised by a strong sequence context, leading to C>T and C>G changes almost solely at TpC sites (21). This contrasts with the weak context dependence observed in the SARS-CoV-2 spectrum. While RNA-editing enzymes may contribute to SARS-CoV-2 mutagenesis, direct damage of cytosine and guanine bases could also be consistent with the observed spectrum. For example, spontaneous cytosine deamination would cause C>U transitions while oxidation of guanine bases could explain G>U transversions (22). Both forms of damage are common mutagenic processes in human cells (23).

Having described the mutational spectrum, we can also derive approximate estimates of the average number of within-host mutations per viral genome in a sample. Since the allele frequency of a mutation represents the fraction of viral RNA molecules in a sample carrying a mutation, we can estimate the mean within-host mutation load by summing the allele frequencies of all observed within-host variants in a sample (12). Importantly, this is an approximation as it assumes that each RNA molecule sequenced, originated from the entire viral genome which is not always the case given the abundance of viral sgRNAs (24, 25). These estimates are also likely to be a conservative lower-bound as they only include mutations at detectable allele fractions in both replicates. Across samples, we found a mean within-host mutation load per viral genome of 0.72 mutations (median 0.37) (Figure 1D). Phylogenetic studies estimate a mutation rate in SARS-CoV-2 of 0.001 mutations/bp/year (26), or 0.082 mutations/genome/day. Thus, the observed within-host mutation load would be consistent with the expected acquisition of mutations during a relatively short span of several days.

To investigate the possible accumulation of *de novo* mutations during the course of an infection, we studied 43 individuals for whom we had multiple samples collected at different timepoints (Figure 3A, Supplementary Figure 4). Overall the number of within-host variants tended to increase over time and this trend was significant using a Poisson mixed model to control for host-specific effects (p=0.007; Figure 3B). To put this finding in context, there was considerable variation in the observed number of within-host variants among samples from the same individual, even if taken on the same day, possibly as a result of bottlenecks caused by the different sampling methods which included sputum, swabs and bronchoalveolar lavage (Supplementary Figure 5). High variability between longitudinal samples has also been observed in six out of nine hospitalised patients in Austria (13). Using data from nine patients with longitudinal samples, the authors found that the divergence in variant frequencies between serial samples from the same patient was on average greater than that between donor-recipient transmission pairs. The authors observed multiple instances of the fixation of a variant over the course of an infection. In contrast, we observed no change in the consensus genome sequence in any of the individuals from whom multiple samples were collected. It is possible that the accumulation of within-host mutations could in future be used to estimate time since infection. However, determining if a consistent signal could be observed given the extensive variation observed between repeated samples on the same day would require analysis of time-series data in a larger number of individuals with high quality epidemiological data.

**Fig. 3.**
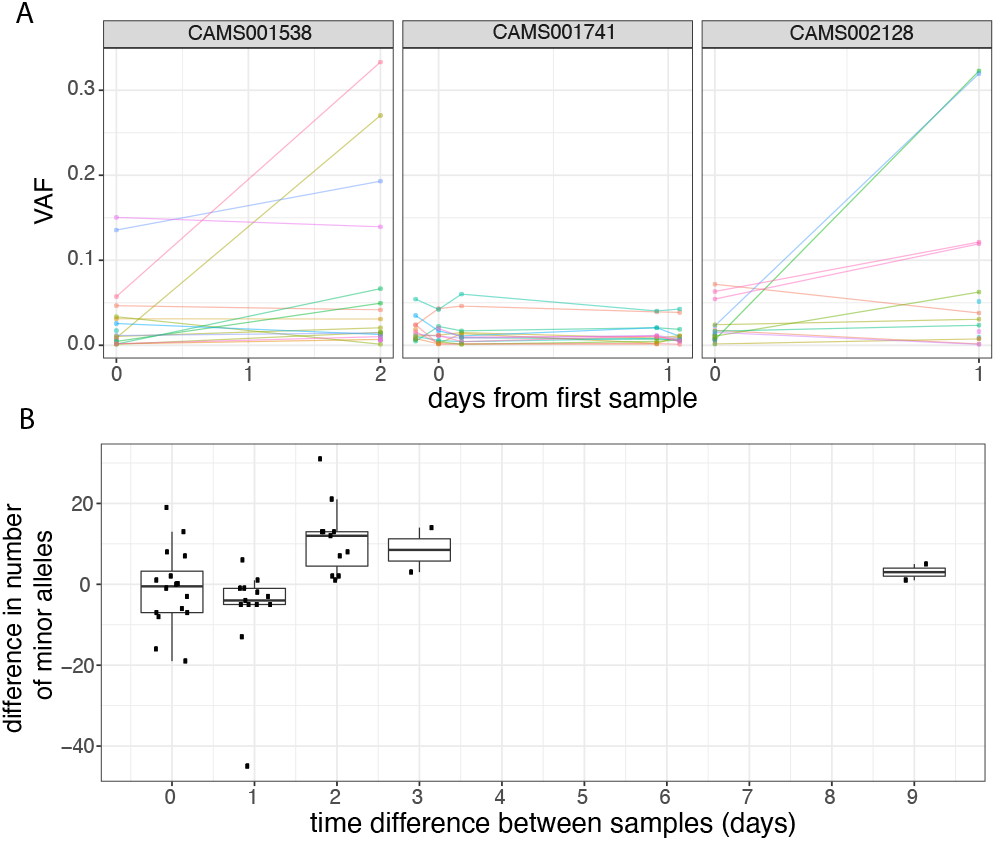
(A) Frequencies of within-host variants for three selected hosts where multiple samples were taken over consecutive days. Samples taken on the same day have been offset by a small distance. Plots for all hosts with multiple samples are given in Supplementary Figure 4 (B) The difference in the number of within-host variants between pairwise combinations of samples taken from the same host.The order for samples taken on the same day was randomised.

### Within-host variants display strong purifying selection against nonsense mutations

To study the extent of selection acting on within-host variants and major variants, we calculated dN/dS ratios using the dNdScv package (Methods). Most commonly-used software to calculate dN/dS use simple substitution models (often using a single transition-transversion ratio), which can lead to considerable biases and false signals of selection under neutrality (28, 29). dNdScv uses a Poisson framework allowing for complex substitution models including context dependence, strand asymmetry, non-equilibrium of substitutions and estimation of dN/dS ratios for missense and nonsense mutations separately. This model is particularly suitable for datasets with low mutation density in lowly- or non-recombining populations, as it is the case in SARS-CoV-2 genomic data (28).

As expected, dN/dS ratios for major variants are under clear purifying selection (Figure 4A), with particularly strong selection against nonsense mutations. Within-host variants reaching moderately high allele frequencies (VAF>10%) display similarly strong purifying selection against missense and nonsense mutations as major variants (Figure 4A), while within-host variants at low allele frequencies appear to be under more relaxed purifying selection, as it may be expected. Purifying selection against nonsense mutations can also be observed at the level of allele frequencies (Figure 4B). Overall, the similarity of dN/dS ratios for major and moderate-VAF within-host variants suggests that selection within hosts, rather than during transmission, may explain a considerable fraction of the extent of purifying selection observed in consensus sequences.

**Fig. 4.**
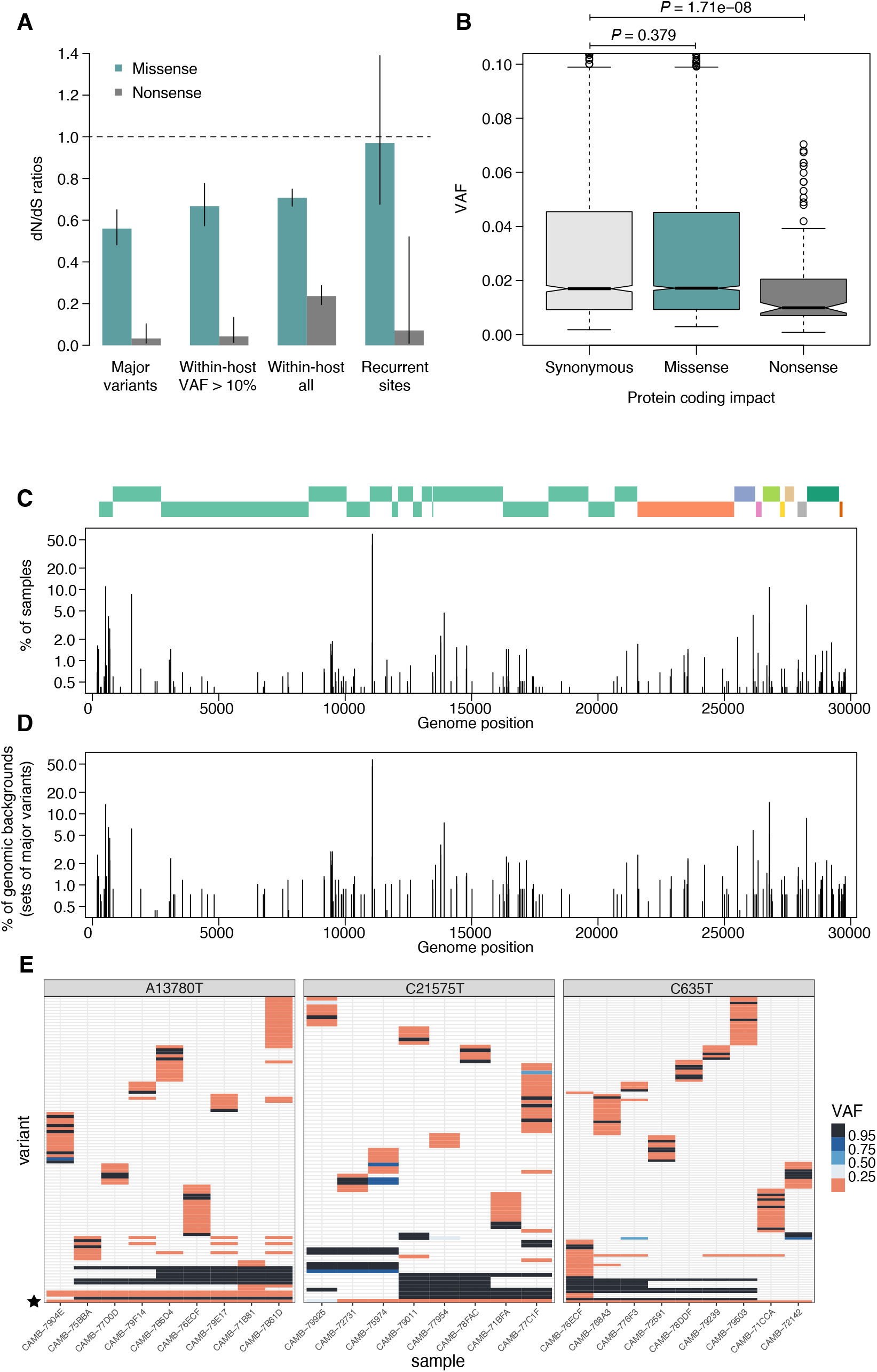
Patterns of selection and recurrent within-host variants. (A) Genome-wide dN/dS ratios for missense and nonsense mutations (Methods). Error bars depict 95% confidence intervals from the Poisson maximum-likelihood model. (B) VAFs of within-host variants as a function of their predicted coding impact. P-values were calculated with Wilcoxon tests. (C) The top panel depicts the coordinates of the annotated peptides in the reference genome, coloured according to their ORF. The bottom panel depicts the frequency at which recurrent within-host variants (defined as those seeing in 5 or more samples) occur in the dataset. (D) Frequency of recurrent within-host variants (as in panel C) across different genomic backgrounds in the dataset (defined as the set of major variants in the sample). (E) Heatmaps of variant allele frequencies in samples containing three common within-host variants are shown. The diversity of major variants (black tiles) across samples is better explained by independent acquisitions of the minority variant rather than transmission

### Many recurrently mutated sites appear to represent mutational hotspots

Some sites across the SARS-CoV2 genome appear to have mutated independently multiple times, resulting in homoplasic sites across the viral phylogeny (30). Some of these sites have also been reported to recurrently appear as within-host variants, although it remains unclear to what extent these events represent recurrent sequencing errors, contamination between samples, coinfection of a sample by multiple lineages, recombination, mutational hotspots or convergent positive selection (3, 13, 30). Coinfection by multiple lineages could possibly arise from superinfection, where an individual acquires infection from multiple sources, or by co-transmission of multiple lineages from host to host following an episode of superinfection.

Figure 4C represents the distribution of recurrent within-host variants across the genome. 190 sites are observed as within-host variants in at least 5 (0.4%) of our samples. Sequencing or PCR errors are unlikely to contribute substantially to the recurrent variants observed in our dataset thanks to the use of replicates and the ShearwaterML algorithm. One mechanism by which the same within-host variant can be observed across multiple closely-related samples is transmission of within-host variants between contacts, when a population of virions is transmitted between hosts. Under this scenario, sharing of within-host variants will be expected between samples with identical major variants, as the preservation of a within-host variant is incompatible with the simultaneous fixation of new major variants (the within-host allele would either be purged or hitchhike and become fixed). Instead, we find that most recurrent within-host variants occur across lineages, independently of the genomic background of major variants present in a sample (Figure 4E). This pattern is suggestive of recurrent mutation, although it could also be consistent with complex histories of superinfection and recombination.

If we accept the hypothesis of recurrent mutation, there remains the question of whether this is caused by mutational hotspots or convergent positive selection. We observed that 25% (51/205) of recurrent (*n* ≥ 5) within-host variants are predicted to cause synonymous mutations and estimates of the dN/dS ratio corrected for the trinucleotide sequence context indicate that, as a group, these recurrent variants are under some purifying selection (Figure 4A). This suggests that most recurrent within-host variants are likely to represent hypermutable sites rather than convergent positive selection. Their mutational spectrum suggests an enrichment for C>U changes, particularly at sites preceded by a pyrimidine, but the mechanisms behind their apparent hypermutability remain unclear (Supplementary Figure 6). Still, this analysis does not rule out the possibility that a minority of recurrent within-host variants are the result of convergent positive selection. A plausible example is the spike glycoprotein mutation D614G, which has rapidly increased in frequency throughout the world: this appears as a within-host variant in 12 of our samples, and typically with a higher allele frequency than other recurrent within-host variants (median VAF 0.56 vs 0.036, Wilcoxon test P=3.98e-4).

### Extensive sharing of minority variants across a diverse genomic background suggests caution is needed when using within-host variants for the inference of transmission

In order to investigate the genetic background of our samples we generated a maximum likelihood phylogeny of all consensus genomes produced by the COG-UK consortium by the end of May 2020, including those for the samples on our dataset (Figure 5A). This showed that our samples represent a broad range of the SARS-CoV-2 genetic diversity found in the UK at that time, and that the diversity observed among our samples was not primarily driven by geographical location.

**Fig. 5.**
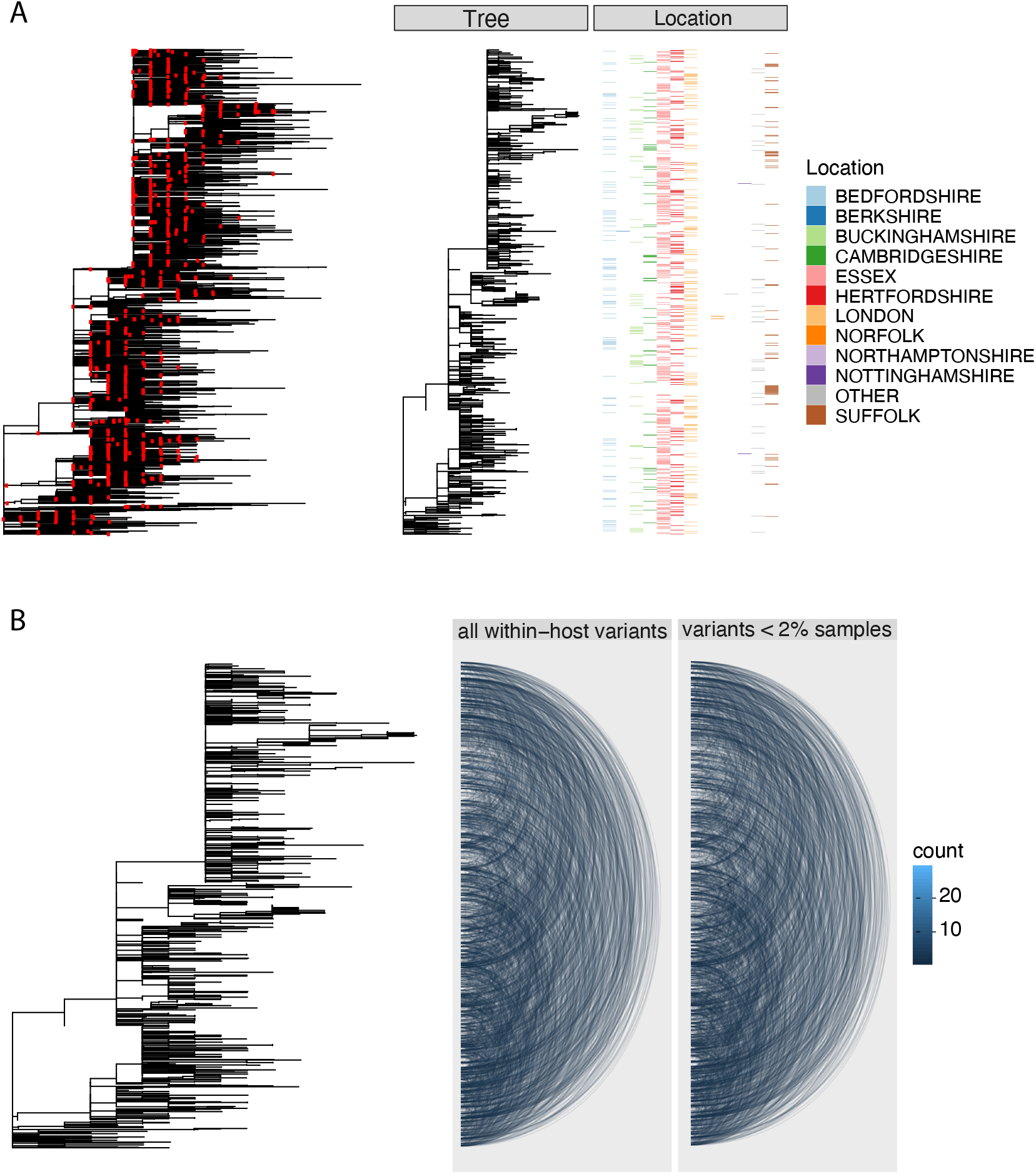
(A) Left panel - a maximum likelihood phylogeny of all COG-UK consensus genomes available on the 29th May 2020. Red dots indicate the location of those samples that were deep sequenced in replicate. Middle panel -the same phylogeny restricted to those samples taken for deep sequencing. Right panel - the region each patient’s home address was located. (B) Left panel - maximum likelihood phylogeny of samples deep sequenced to investigate minority variants. Middle panel - links are drawn between tips of the phylogeny that share within-host minority. Right panel - links restricted to those variants seen in less than 2% of individuals.

To explore the relationship between within-host variants and the consensus phylogeny, we identified within-host variants that are shared between samples, as illustrated by links between the tips of the phylogenetic tree in Figure 5B. This confirmed that within-host variants are often shared between samples that are distant on the consensus phylogeny. Both a high level of recombination and a large transmission bottleneck would be required for within-host variants to be maintained across long transmission chains in order to explain the simultaneous preservation of some within-host variants with the fixation of others. While, recent studies have suggested that the bottleneck in SARS-CoV-2 transmission can be on the order of 10^2^-10^3^ virions, they also identified substantial overlaps in the fraction of shared minority variants between samples unrelated by close transmission (13, 31). The sharing of within-host variants between consensus genomes as divergent as 10 SNPs, indicating multiple months of separation between the samples suggests that a more likely explanation is that many of these shared within-host variants are the result of recurrent mutation or co-infection (Fig 6D).

**Fig. 6.**
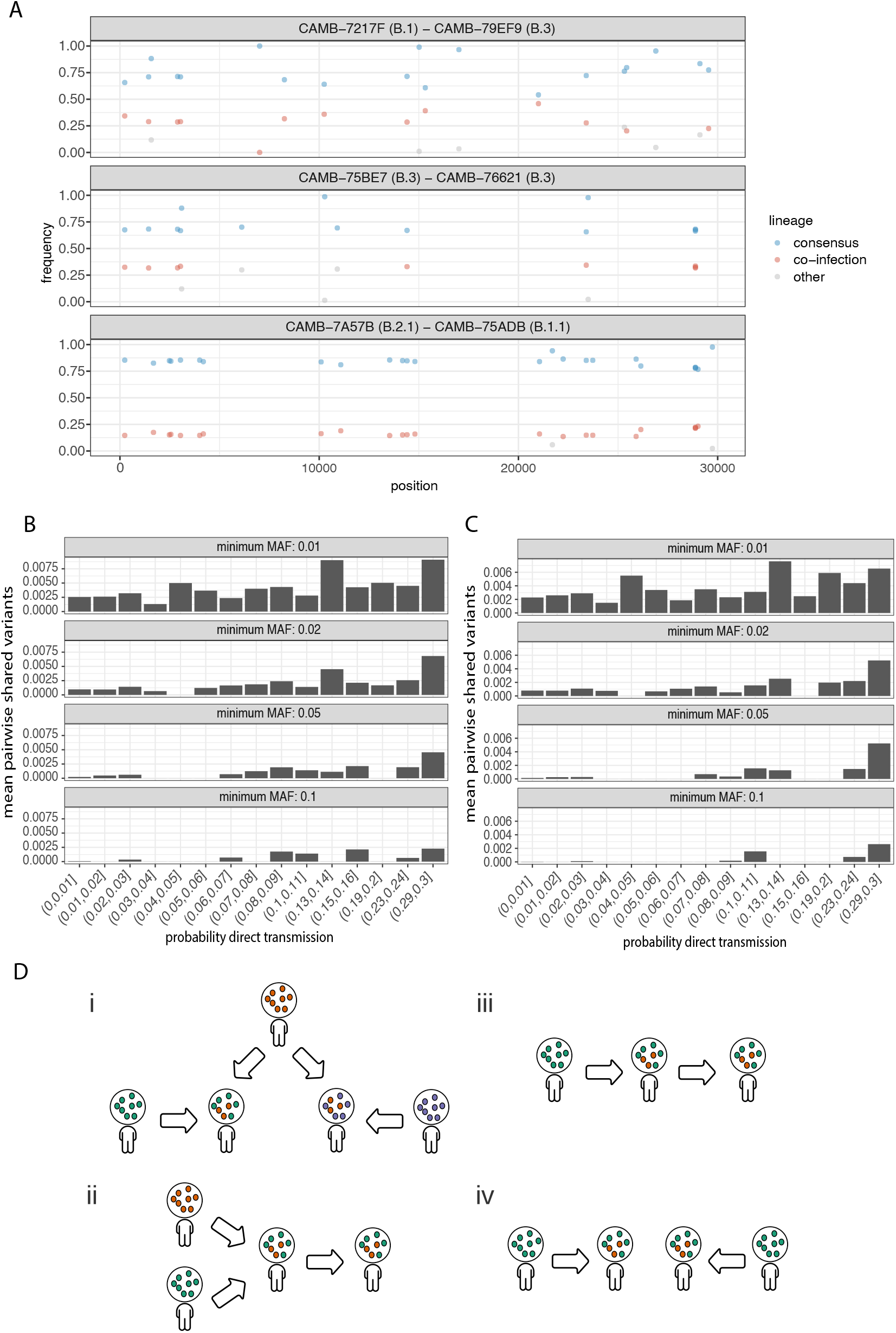
(A) An example of three samples identified as potential mixtures. The consensus lineage is given first and coloured blue while the potentially co-infecting lineage is given second and coloured red. Minority variants that do not match the co-infecting lineage are coloured grey. (B) The median number of shared iSNVs shared by each pair of samples binned by the probability they were the result of a direct transmission according to the model of Stimson et al., 2019. Results, with a minimum minor allele frequency of 0.01, 0.02, 0.05 and 0.1 are shown in each of the facets. (C) The same plot as 3B but having removed all samples that were inferred to be mixed infections. (D) A diagram demonstrating the four scenarios that can lead to shared within-host variants. (i) Super-infection of a common strain. (ii) Super-infection followed by co-transmission (iii) Transmission of the within-host variants through a large bottleneck. (iv) Independent *de novo* acquisitions of the same within-host variants. Shared within-host variants in scenarios (ii, iii) are concordant with the transmission tree while (i, iv) are discordant, potentially confounding transmission inference efforts.

To further investigate the correspondence between shared within-host variants and transmission, we used the transcluster algorithm to infer the probability distribution of the number of intermediate hosts that separate each pair of samples(32). This pairwise approach accounts for the serial interval of the virus as well as its evolutionary rate using the difference in the number of SNPs between each pair of consensus genomes. Fig 6B illustrates the relationship between the mean number of shared within-host variants and the probability that the respective infections were a result of direct transmission. A weak correlation with transmission probability was found for shared within-host variants of high allele frequency, but not for those of low allele frequency.

### Co-infections correlate with the consensus phylogeny consistent with cases of superinfection induced by population structure

It has been proposed that coinfection by different lineages is a significant source of within-host variation in SARS-CoV-2 (3). Co-infection by multiple lineages would result in the allele frequencies of divergent bases to appear as within-host variants and reflect the proportions of either haplotype. Detecting instances of co-infection is important in the context of both transmission and to accurately characterise the within-host mutation rate (9, 33). To identify possible co-infected samples we compared the site frequency spectrum for each of our samples with all of the consensus genomes in the COG-UK dataset at the end of May 2020. A linear model was used to identify mixtures of two consensus genomes that could explain the allele frequencies of variants within a sample better than a single consensus genome. Additional mixtures were considered if two consensus genomes could explain at least two additional within-host variants over that of a single consensus genome. In this case the additional consensus genome was restricted to have at most one variant that was not found in the sample. Samples identified by the model as being a possible mixture of consensus genomes were then visually inspected to determine whether the putative co-infections were convincing. This resulted in 36 putative co-infected samples, with a representative example shown in Figure 6A. The frequencies of within-host variants of all 36 samples, along with consensus genomes that comprise the putative mixture, are shown in Supplementary Figure 7. To determine the potential impact of co-infections on transmission inference we excluded all 36 samples and re-ran the pairwise transmission analysis using the transcluster algorithm on the remaining samples. Figure 6C indicates that the remaining within-host variants no longer correlate as well with the probability of direct transmission, suggesting that much of the correlation observed previously was potentially driven by co-infections.

It has been suggested that correlations between co-infections and the consensus phylogeny could be driven by the cotransmission of multiple strains (3). However, as the transmission signal in the minority variants in the remaining samples has reduced; a more likely explanation is that coinfection of certain strains is driven by multiple episodes of superinfection (co-infection from two different infection sources). As these samples and those from the Lythgoe study were acquired from hospitalised patients, the correspondence with the consensus phylogeny could then be driven by structured superinfection within hospital COVID-19 wards. The prevalence of infection within hospitals, particularly during the first ‘wave’ of the COVID-19 pandemic was often considerably higher than in the community with up to 50% of the consultant A&E workforce at one Welsh hospital testing positive in April, 2020 (34). This contrasts with the finding that approximately 6% of the UK population had been infected with SARS-CoV-2 by the end of June, 2020 (35). Within hospital transmission was also common in the early stages of the outbreak in Wuhan with 41% of 138 patients were found to have contracted SARS-CoV-2 in hospital (36). While it is not possible to rule out cross-contamination between samples, this contamination would have to be structured to explain the correlations between the inferred co-infections and the consensus phylogeny. Batch effects were carefully controlled for by Lythgoe et al., and as we identified a similar signal using an independent dataset with samples sequenced in replicate, structured superinfection provides a better explanation. In cases of higher prevalence, repeated episodes of superinfection rather than co-transmission could complicate the use of within-host minority variants in transmission inference (Figure 6D).

## Discussion

We find a considerable amount of genetic diversity within individual SARS-CoV-2 infections that cannot be explained by technical artefacts. This is consistent with the quasispecies population structure typically found in RNA viruses (5, 37), including related betacoranaviruses SARS-CoV-1 and MERS (38, 39). By analysing the frequency of variants within individual samples, we estimate that each viral genome has an average of 0.72 variants relative to the consensus genome sequence for that sample. Since most of these samples were probably collected more than a week after the time of infection, and since the SARS-CoV-2 genome is known to acquire approximately one mutation every two weeks, these findings are broadly consistent with the hypothesis that within-host variation is largely due to the accumulation of *de novo* mutations within the host. Further support for this hypothesis comes from our analysis of hosts sampled at multiple timepoints, showing that the number of within-host variants tends to increase during the course of an infection. Increased numbers of within-host variants have also been observed in immunocompromised patients: consistent with the acquisition of *de novo* mutations within the host (40).

These data show that within-host variants have a similar mutational spectrum to major variants that define between-host variation. Both are characterised by clear purifying selection, as would be expected if virions with disadvantageous mutations failed to survive or propagate within the host. Strikingly, the mutational spectrum of SARS-CoV-2 exhibits a near complete asymmetry between the plus and minus strands, and is dominated by C>U and G>U mutations. This seems consistent with RNA-damage or RNA-editing of the plus strand. Whilst mutagenesis by APOBEC enzymes could play a role, the spectrum we observe is very different to that seen in human cells. Instead, direct damage of cytosine and guanine bases could also be consistent with the observed spectrum. Many of the within-host variants that we have identified a re s hared b etween i nfected i ndividuals located on distant branches of the consensus phylogeny, and this is congruent with previous reports that the SARS-CoV-2 consensus phylogeny has many homoplasies that also appear as within-host variants (3). It appears likely that this is largely due to recurrent mutation although co-infection between lineages could also be a relevant factor in high transmission settings. The vast majority of our samples appear to comprise a single lineage but we find evidence of putative co-infection by multiple lineages in approximately 3-4% of samples, which is likely an underestimate of the extent of co-infection in our dataset.

In other viral and bacterial diseases, within-host variants provide a valuable source of information for the inference of transmission chains (33, 41–43) and have been shown to improve the accuracy of inferences based on consensus genomes (44–47). In the case of SARS-CoV-2, Lythgoe and colleagues have reported a geographical structure to the sharing of within-host variants, and have proposed that this might be explained by localised episodes of superinfection (where an individual acquires infection from multiple sources) and subsequent co-transmission of multiple lineages and their recombinants through the local host population (3). We find some evidence of a correlation between sharing of within-host variants and the probability of direct transmission, a signal that we find is partially driven by co-infection, but the current dataset lacks the epidemiological resolution to evaluate the utility of this in reconstructing transmission chains, which might depend on the prevalence of infection and other local circumstances such as superinfection within COVID-19 hospital wards.

In summary, these data show that much of the genetic diversity of the SARS-CoV-2 viral population resides within individual hosts. Within-host diversity is generated by the accumulation of *de novo* mutations during the course of infection but can also result from co-infection with different lineages. Analysis of within-host variation could potentially be used to improve the inference of transmission chains, but this approach requires caution because of the confounding effects of recurrent mutation and unrelated episodes of superinfection. More detailed studies are required to evaluate the transmission bottlenecks that govern the propagation of within-host variants from host to host, and to examine the patterns of within-host variation associated with different epidemiological circumstances.

## Methods

### Sample selection and ethics

The 1,181 samples were taken as a random subset from a larger prospective study into SARS-CoV-2 infections at Cambridge University Hospitals National Health Service Foundation Trust (CUH; Cambridge, UK), a secondary care provider and tertiary referral centre in the East of England (4, 48). This study was done as part of surveillance for COVID-19 under the auspices of Section 251 of the National Health Service Act 2006. It therefore did not require individual patient consent or ethical approval. The COVID-19 Genomics UK (COG-UK) study protocol was approved by the Public Health England Research Ethics Governance Group.

### Variant calling

Variant calling was performed using ShearwaterML, which is available as part of the deepSNV R package (11, 12). In order to create a base-specific error model for the SARS-CoV-2 genome, we randomly selected sequencing data from 100 samples (50 from each set of duplicates) from the 468 samples that met the following criteria: (1) *ρ* value from beta-binomial model ≤ 0.02; (2) proportion of genome with at least 500× coverage ≥0.9 for both duplicates; (3) absolute difference in median coverage between replicates ≤20,000; and (4) only one sample sequenced from a given donor. For each sample, non-reference sites with VAF ≥ 0.01 were set as uninformative (i.e. depth for all base types were set to 0) in order to enable calling of recurrent, high VAF mutations. However, two sites (11074 and 25202) were effectively excluded from variant calling as they were set as uninformative for all samples in the normal panel.

Variants were called separately for each set of duplicates. The initial ShearwaterML calls were filtered using the following criteria: (1) total depth at variant site *ge*100; and (2) Benjamini-Hochberg False Discovery Rate q-value ≤0.05 in one duplicate and unadjusted p-value ≤0.01 in the other duplicate. The adjustment of p-values was performed by considering the five mutation types (three non-reference bases, insertions and deletions) at all sites with ≥100× coverage in the 1,181 samples from a given duplicate set. Variants at consecutive sites were merged into single events if the difference in VAFs ≤ 0.05.

### Beta-binomial modelling of replicate samples

To quantify the level of discordance in the allele frequencies between technical replicates, for each pair replicates we first identified a set of variable sites, which are expected to contain both artefacts and within-host mutations. All non-reference variable sites with coverage in both replicates *>* 100x and mean VAF from both replicates higher than 1% and lower than 90% were considered for the beta-binomial modelling. In Illumina sequencing protocols where adapters are ligated before amplification and PCR duplicates can be removed computationally, VAFs may be expected to show binomial variation between technical replicates. However, amplicon sequencing can lead to preferential amplification of some molecules leading to higher variation in VAFs. We quantified the extent of variation above binomial sampling in the VAFs of variable sites between replicates by fitting a beta-binomial model to the allele counts of variable sites, obtaining a maximum likelihood estimate of the *ρ* (overdispersion) parameter for each pair of replicate samples. Let *x*_1*,j*_ and *x*_2*,j*_ be the alternative (non-reference) allele counts of variable site *j* in replicates 1 and 2 of a given sample, and *n*_1*,j*_ and *n*_2*,j*_ be the local coverages at the site. We can calculate an approximate likelihood for the beta-binomial model using the following equation, where *P* depicts the beta-binomial density function:

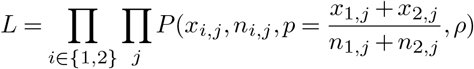

A maximum likelihood estimate for the overdispersion parameter was obtained by grid search. The overdispersion parameter, measuring the level of discordance in allele frequencies between the replicates, was found to correlate strongly with the Ct value of the sample.

### Mutational spectrum

The mutational spectra shown in Figure 1 were generated assuming that the allele in the reference sequence (SARS-CoV-2 isolate Wuhan-Hu-1, MN908947.3) represents the ancestral allele. The normalised mutation rates (r) for each of the 192 possible changes (j) in a trinucleotide context were calculated as:

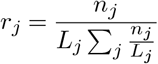

Where *n_j_* is the total number of mutations observed for a trinucleotide change *j*, and *L_j_* is the total number of times that the corresponding trinucleotide is present in the reference genome (MN908947.3). When observing the same mutation in multiple samples, it is not always straightforward to determine the number of independent mutational events that this represents. For major variants, this can be better estimated using a phylogeny, although recurrent hotspots can cause errors in the phylogenetic reconstruction. For simplicity, mutations observed across samples as major variants were counted only once for the estimation of *n_j_*. For within-host variants, a more relaxed approach was used since, based on our results, multiple instances of the same within-host variant across samples are more likely to represent independent events. Within-host variants observed in more than 5 samples were only counted 5 times for the calculation of *n_j_*, to avoid a small number of highly-recurrent hotspots from dominating the trinucleotide spectrum.

### Selection analysis

Analyses of selection were carried out using the dNdScv package (28). In contrast to traditional implementations of dN/dS developed for divergent sequences, which rely on Markov-chain codon substitution models (49), dNdScv was developed for comparisons of closely related genomes. Under low divergence rates, such as those in the densely sampled SARS-CoV-2 phylogeny, observed changes typically represent individual mutational events, which can be modelled using a Poisson framework instead of Markovchain substitution models. This enables the use of more complex and realistic substitution models, including context dependence, strand asymmetry, non-equilibrium sequence composition and separate estimation of dN/dS ratios for missense and nonsense mutations. These changes can be important to avoid false signals of negative or positive selection under neutrality when using simplistic substitution models (28). Here we used the default substitution model in dNdScv, which uses 192 rate parameters to model all possible mutations in both strands separately in a trinucleotide context, as well as two *ω* parameters to estimate dN/dS ratios for missense and nonsense mutations separately. dNdScv files and code needed to generate dN/dS ratios from SARS-CoV-2 data are available as Supplementary Code.

dN/dS ratios on polymorphism data need to be interpreted with caution. This is both because dN/dS ratios can be time dependent (50), providing weaker signals of selection for more recent changes, and because dN/dS ratios can behave non-monotonically with respect to selection coefficients under idealised conditions of free recombination when using nucleotide diversity within a population (51). The latter can result from strong positive selection causing a reduction in the effective population size (and in nucleotide diversity) of non-synonymous sites under free recombination. However, the potential loss of monotonicity should not be a concern in our analyses of SARS-CoV-2 data. This is both because free recombination between adjacent sites is unrealistic for SARS-CoV-2 and because collections of mutations were identified by comparison of derived sequences to an ancestral reference genome, instead of by comparison of two derived sequences at any one time.

### Consensus phylogeny construction

Consensus genomes for each replicate were generated using the ARCTIC SARS-CoV-2 bioinformatics pipeline. A multiple sequence alignment was generated using MAFFTv7.464 and any sites that were discordant between replicates were set to be ambiguous (52). Sites that have previously been identified to create difficulties in generating phylogenies were masked using bedtools v2.29.2 using the VCF described in De Maio et al, 2020 (30, 53). Fasttree v2.1.11 was used to generate a maximum likelihood phylogeny (54).

### Transmission model

To investigate transmission, samples were only considered if both replicates produced high quality consensus genomes. When multiple samples from the same host were available the earliest sample was used. Pairwise SNP distances were generated between the consensus genomes using pairsnp v0.2.0 (55). The distribution of the underlying number of intermediate transmission events between each pair of samples was then inferred using an implementation of the transcluster algorithm (32, 56). The serial interval and evolutionary rate were set to 5 days and 1e-3 substitutions/site/year (26, 57).

### Identification of potential mixed infections

Potential mixed infections were identified using a linear model by testing whether the allele frequencies in a sample could be better explained by the inclusion of an additional consensus genome from the COG-UK dataset of the 29th May 2020. Additional samples mixtures were considered if the addition of a COGUK consensus genome could explain at least 2 iSNVs and have at most 1 variant that was not found in the alleles of the sample. This identified 54 putative mixtures which were then screened manually to obtain 36 potentially mixed samples. The code used to run this analysis is available in the supplementary materials.

## SOFTWARE AVAILABILITY

Analysis code available from: https://github.com/gtonkinhill/SC2_withinhost

## DATA AVAILABILITY

Accession numbers for the sequence data is given in Supplementary Table 3

## ACKNOWLEDGEMENTS

This work was funded by COG-UK, supported by funding from the Medical Research Council (MRC) part of UK Research & Innovation (UKRI), the National Institute of Health Research (NIHR) and Genome Research Limited, operating as the Wellcome Sanger Institute; the Wellcome Trust (Senior Fellowship to IG ref: 207498/Z/17/Z and PhD Scholarship to GTH ref: 204016/Z/16/Z); the Academy of Medical Sciences & the Health Foundation (Clinician Scientist Fellowship to MET; and the Cambridge NIHR Biomedical Research Centre (MET).

## Supplementary Figures

**Supplementary Figure 1:**
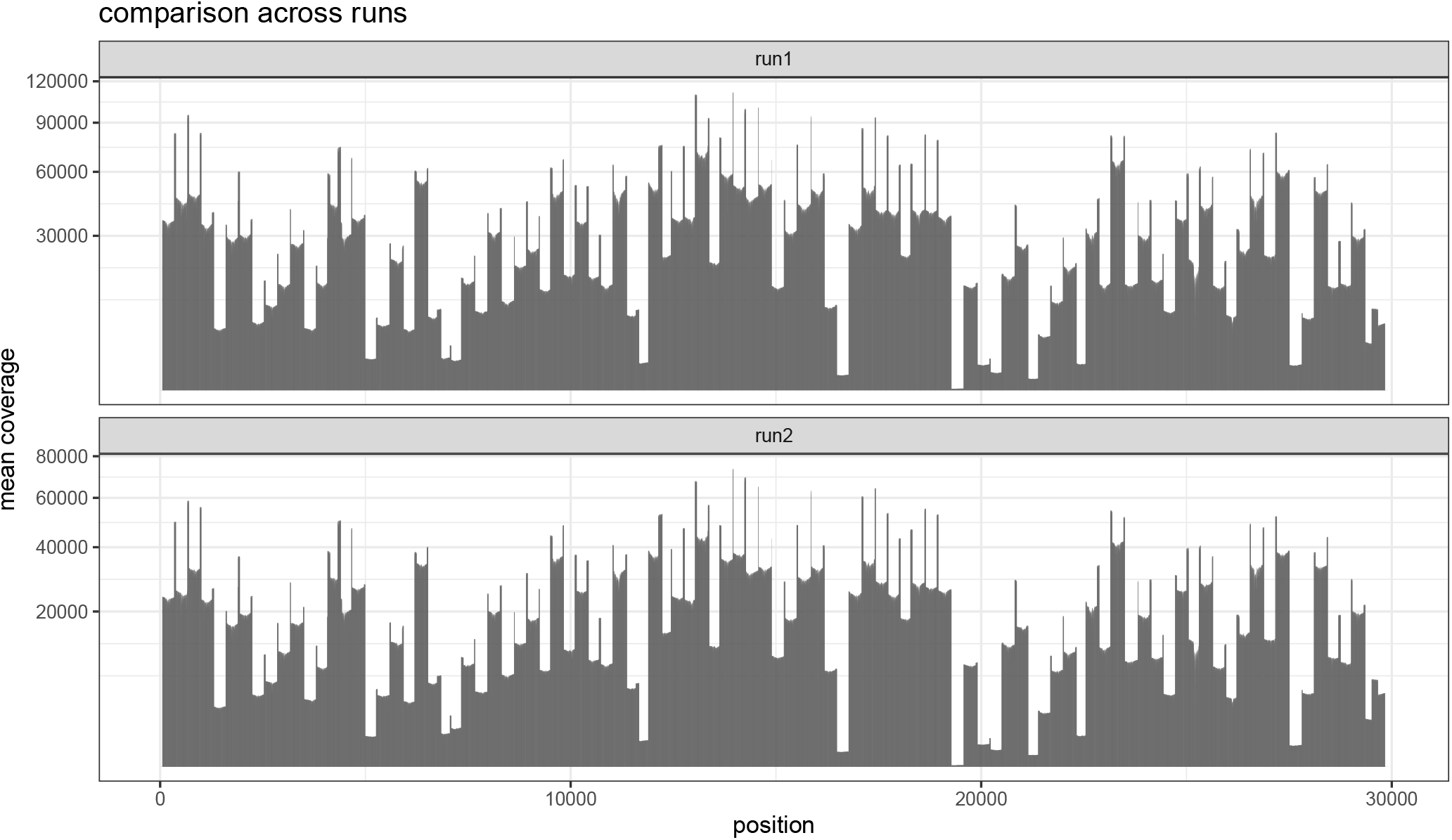
Barplots indicating the mean sequencing depth across the SARS-CoV-2 reference genome for the two replicate runs of the 1,181 samples.

**Supplementary Figure 2:**
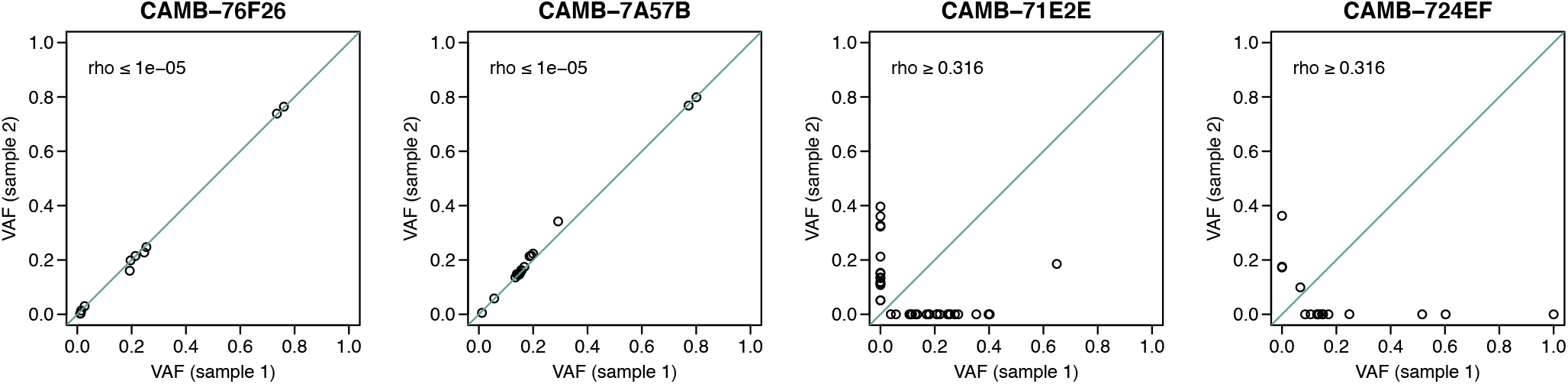
Dot plots indicating the concordance between variant allele frequency estimates across sequencing replicates in four samples.

**Supplementary Figure 3:**
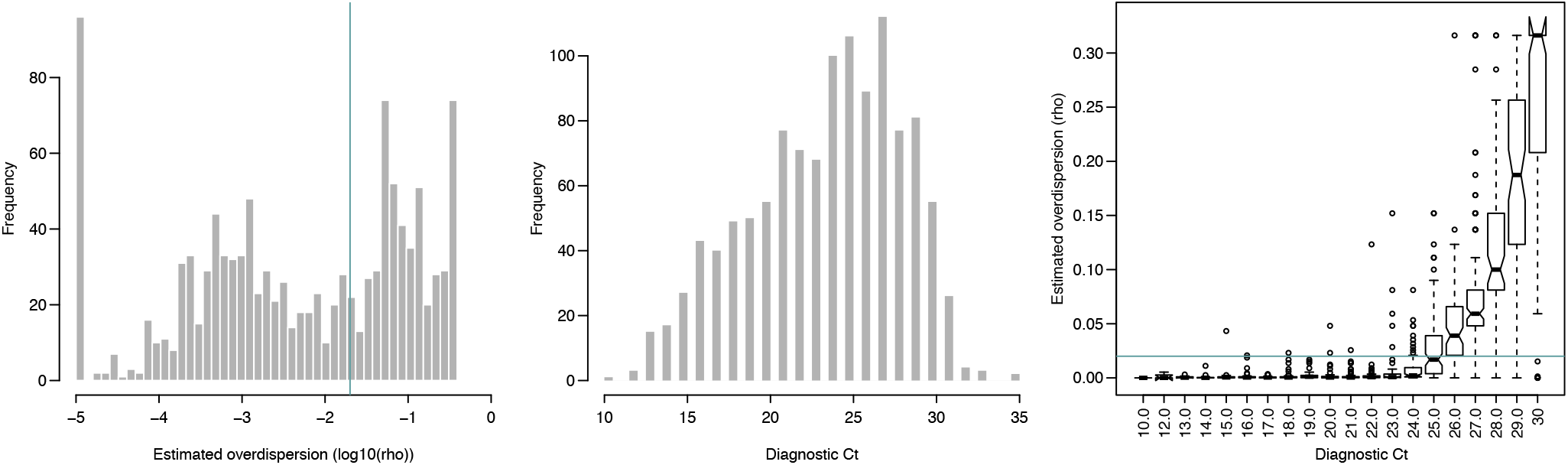
(A) Histogram of estimated log10(*ρ*) values. Green line represents *ρ* = 0.02 in (A) and (C), as a suggested acceptable level of discordance between replicates. 58% of all samples in the cohort had *rho ≤* 0.02. (B) Histogram of Ct values in the cohort. (C) Estimated *ρ* value as a function of Ct.

**Supplementary Figure 4:**
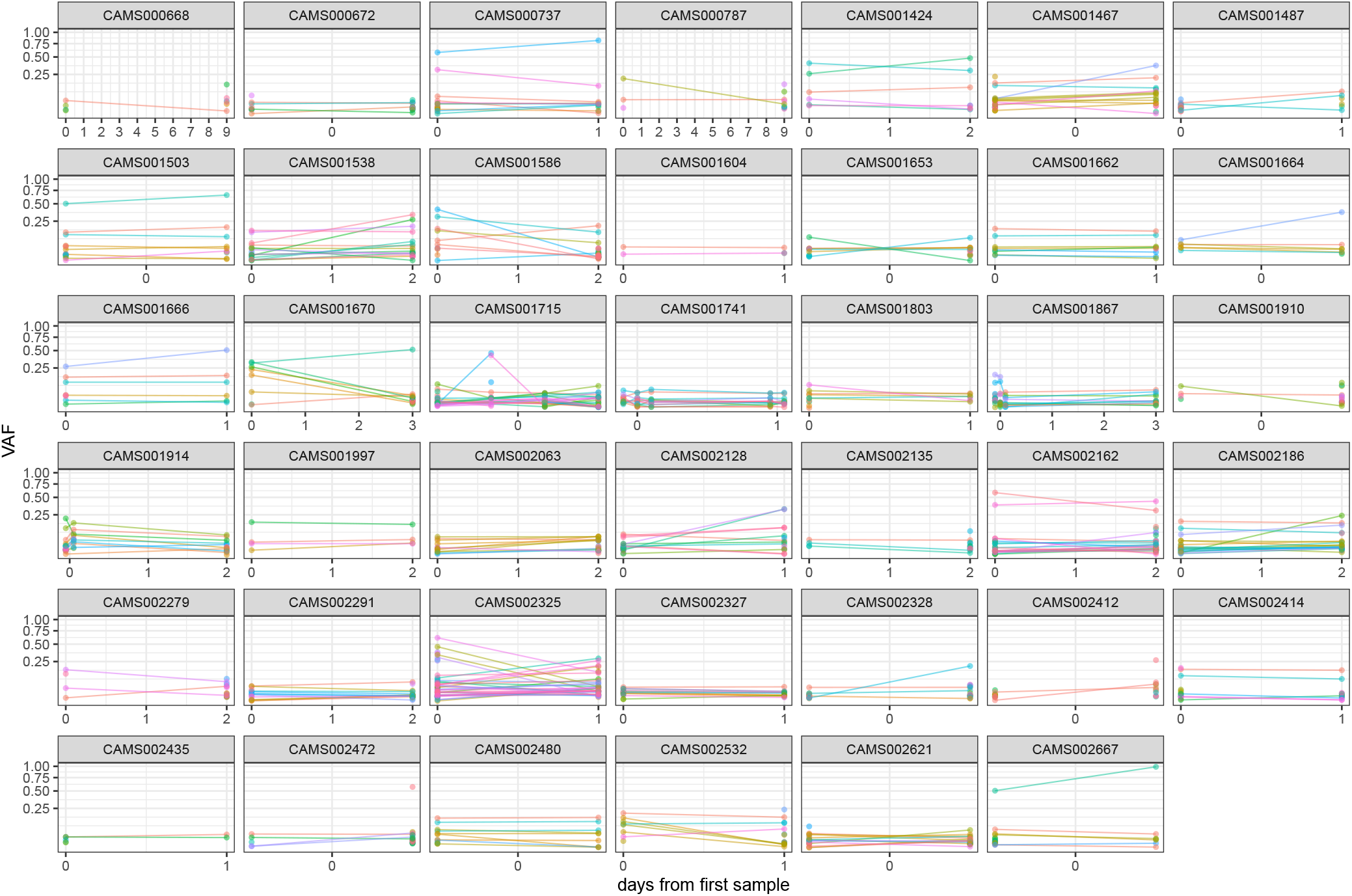
Frequencies of within-host variants for all hosts where multiple samples were taken over consecutive days. Samples taken on the same day have been offset by a small distance to allow for comparison.

**Supplementary Figure 5:**
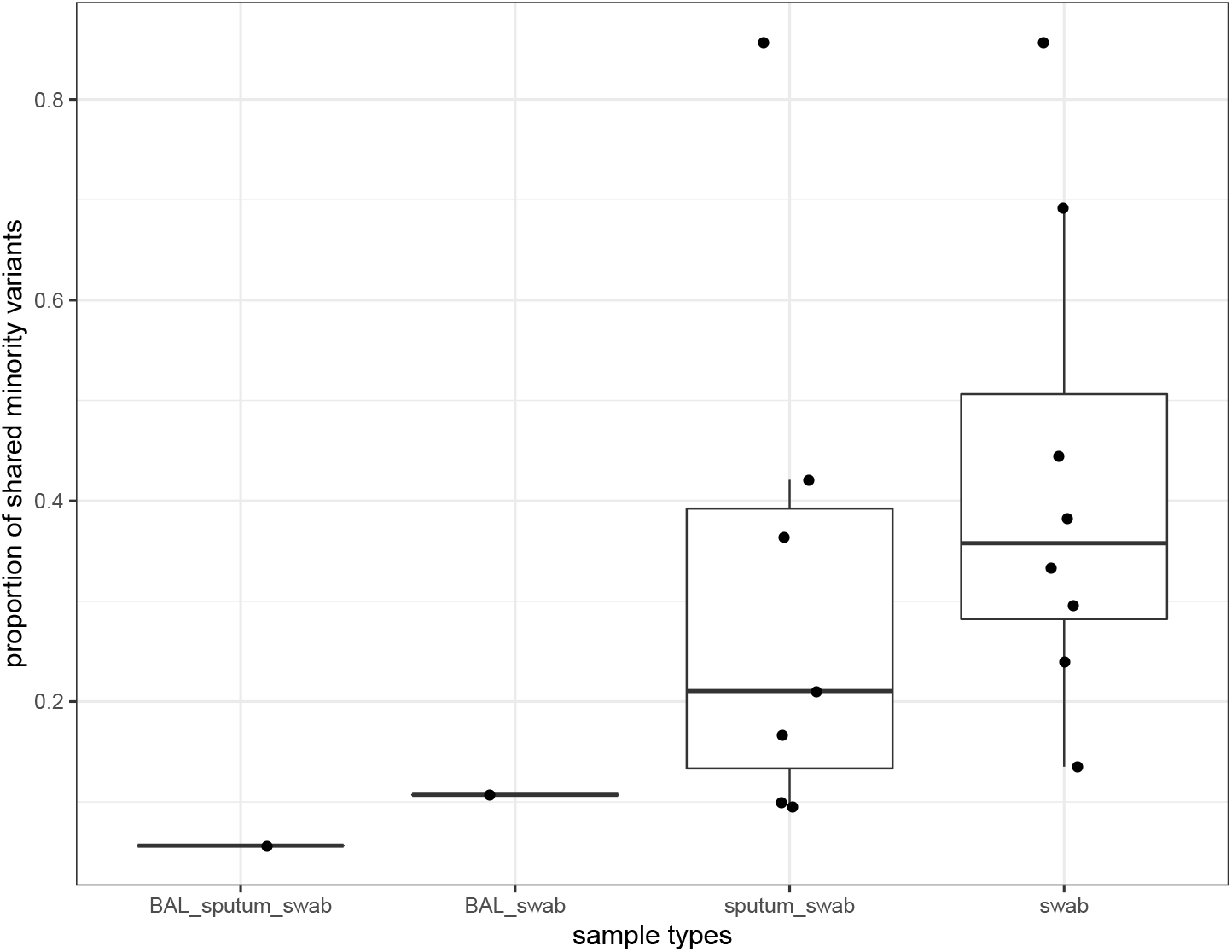
Proportion of shared variants between each pair of samples taken from the same host on the same day. Pairs are split by sampling method which included sputum, swabs and bronchoalveolar lavage.

**Supplementary Figure 6:**
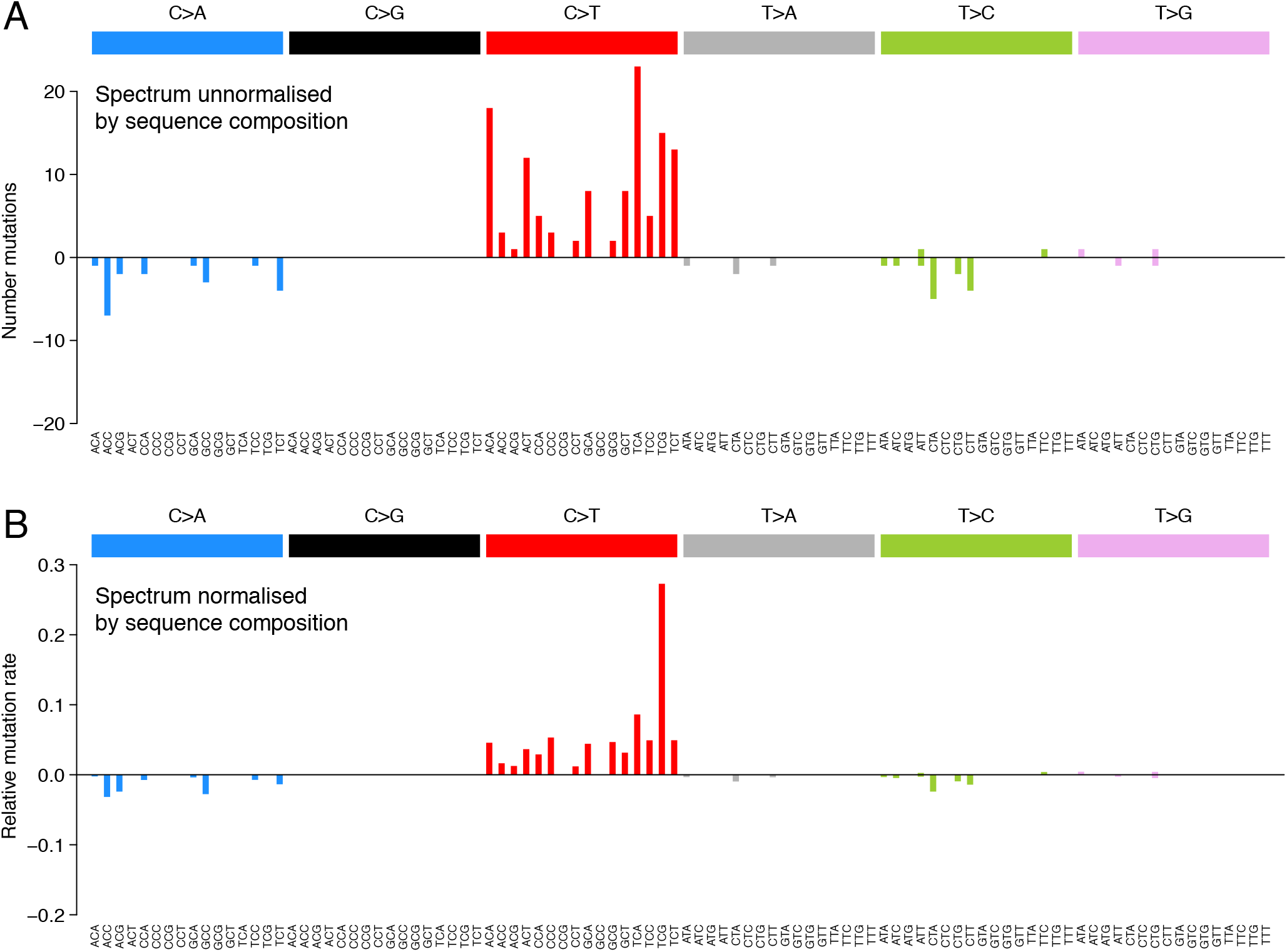
The mutational spectra in a 96-trinucleotide context of recurrent within-host variants.

**Supplementary Figure 8:**
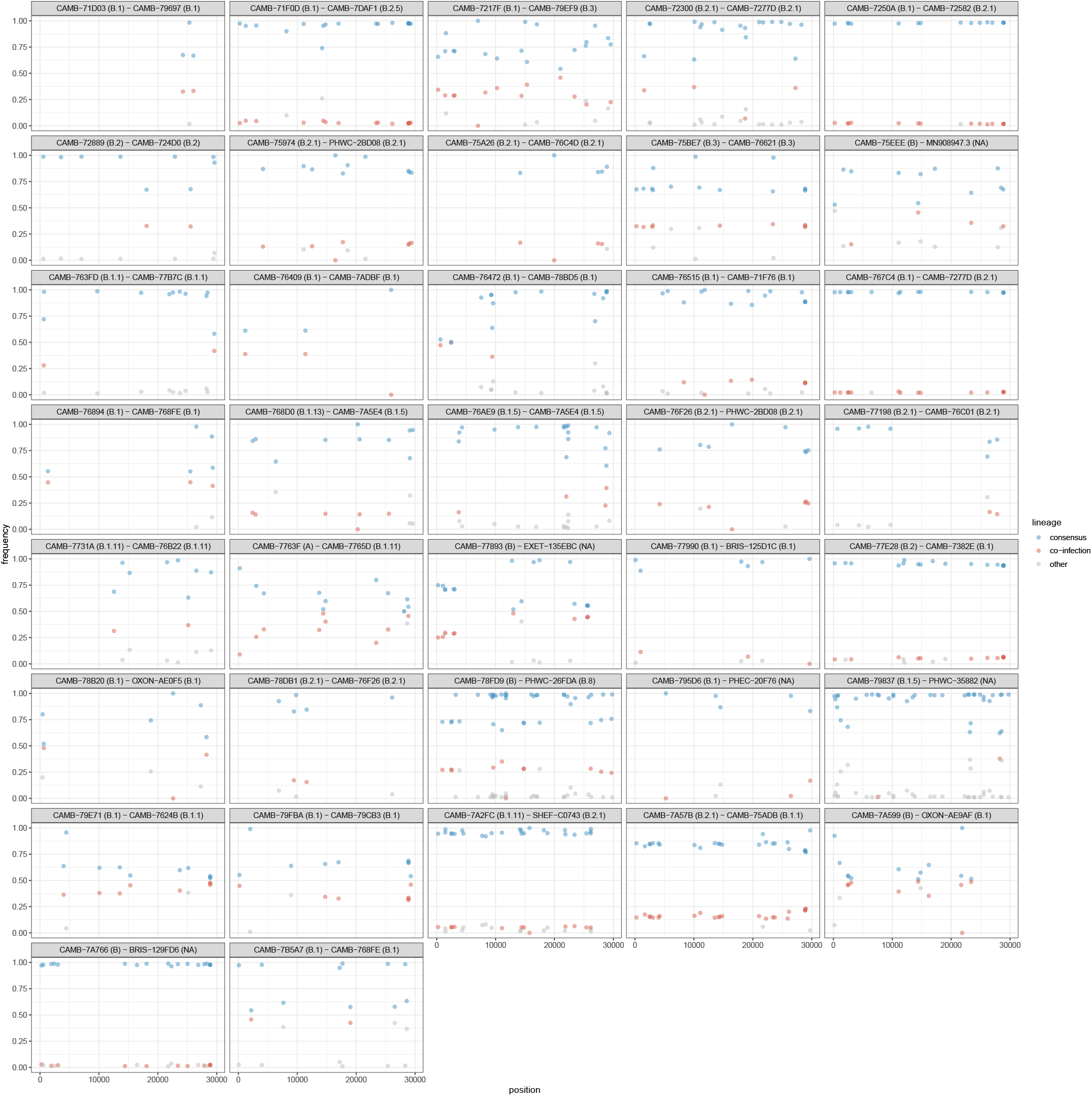
All samples identified as potential mixtures. The consensus lineage is given first and coloured blue while the potentially co-infecting lineage is given second and coloured red. Minority variants that do not match the co-infecting lineage are coloured grey.

## Notes

https://github.com/gtonkinhill/SC2_withinhost

### Competing Interest Statement

The authors have declared no competing interest.

